# Dissociation of direct and peripheral transcranial magnetic stimulation effects in nonhuman primates

**DOI:** 10.1101/2022.12.26.521973

**Authors:** Nipun D Perera, Ivan Alekseichuk, Sina Shirinpour, Miles Wischnewski, Gary Linn, Kurt Masiello, Brent Butler, Brian E Russ, Charles E Schroeder, Arnaud Falchier, Alexander Opitz

**Affiliations:** Department of Biomedical Engineering, University of Minnesota, Minneapolis, MN, USA; Translational Neuroscience Lab Division, Center for Biomedical Imaging and Neuromodulation, The Nathan S. Kline Institute for Psychiatric Research, Orangeburg, NY, USA; Departments of Psychiatry and Neurosurgery, Columbia University College of Physicians and Surgeons, New York City, NY, USA; Department of Psychiatry, NYU Grossman School of Medicine, New York City, NY, USA

**Keywords:** Transcranial magnetic stimulation, local field potentials, non-human primates, invasive electrophysiology

## Abstract

Transcranial magnetic stimulation (TMS) is a non-invasive brain stimulation method that is rapidly growing in popularity for studying causal brain-behavior relationships. However, its dose-dependent direct neural mechanisms, i.e., due to electric field or connectivity, and peripheral sensory co-stimulation effects remain debated. Understanding how TMS stimulation parameters affect brain responses is vital for the rational design of TMS protocols. Studying these mechanisms in humans is challenging due to the limited spatiotemporal resolution of available non-invasive neuroimaging methods. Here, we leverage invasive recordings of local field potentials in non-human primates to study TMS mesoscale responses. We demonstrate that early TMS-evoked potentials show a sigmoidal dose-response with stimulation intensity. We further show that stimulation responses are spatially specific. We employ several control conditions to dissociate direct neural responses from auditory and somatosensory co-activation. These results provide crucial evidence regarding TMS neural effects at the brain circuit level. Our findings are highly relevant for interpreting human TMS studies and biomarker developments for TMS target engagement in clinical applications.

## INTRODUCTION

Modulating local neural activity has great potential to restore affected brain regions in neurological and psychiatric disorders. Non-invasive neuromodulation is a cost-effective tool for large-scale use in patients with a wide range of symptoms and symptom severity. The current gold standard for non-invasive neuromodulation in clinical and basic research is transcranial magnetic stimulation (TMS). TMS can induce action potentials in neurons and, through its repeated application, brain plasticity [1,2]. TMS-based therapies received FDA approval to treat major depressive disorder, obsessive-compulsive disorder, and nicotine addiction [3,4]. Despite its increasing use in research and clinical applications, the physiological effects of TMS are not fully understood. This gap of knowledge has hampered rational dose selection and optimization of TMS parameters. In particular, a clear physiological marker of TMS target engagement would be crucial to advance clinical applications in brain regions other than the motor cortex, such as the dorsolateral prefrontal cortex (DLPFC).

In human research, TMS physiological effects can be measured non-invasively through motor-evoked potentials [5–7], functional magnetic resonance imaging [8–10], and electroencephalography (EEG) [11–16]. Combined TMS-EEG is a promising method to study neural population responses to TMS with high temporal accuracy. A primary way of investigating transient stimulation effects on EEG is by measuring TMS-evoked potentials (TEPs). For example, causal brain dynamics during TMS perturbation in motor, visual, and prefrontal brain regions [11,14] and modulation of brain state during behavior [12,15] have been studied through TMS-EEG. It has also been used to quantify cortical inhibition paradigms that are resulting from TMS perturbation [13,16]. However, despite its promise to study the neural responses of TMS, the interpretation of human TMS-EEG neurophysiological results is challenging.

One challenge for the interpretation of TMS-EEG is a poor spatial localization of results due to volume conduction, meaning that the location of TEP sources is unclear. Second, multisensory side effects during TMS include auditory and somatosensory co-activation (by TMS click sound and muscle and peripheral nerve co-stimulation), which also affect brain responses. This result in a challenge to disentangle direct brain stimulation effects arising from electric field or connectivity and peripheral sensory stimulation effects [17,18]. To systematically investigate TMS physiological mechanisms, several control conditions carefully accounting for various confounds are required [19,20]. Non-human primate (NHP) models can overcome the main limitations of human TMS-EEG. Using invasive electrophysiological recordings in NHPs offers a unique opportunity to study the physiological effects of TMS with high spatiotemporal precision and control for somatosensory effects. Further, compared to smaller animal models, the size and cortical folding patterns of the NHP brain will result in comparable TMS-induced electric fields as in the human brain, making them an ideal translational model [21].

Previous work using TMS in NHPs has studied the effect of single-pulse TMS on single-unit activity [22,23]. These findings have shown that TMS (compared to sham stimulation) causes neural activity within milliseconds, and this effect is observed in various neuron types [22]. Furthermore, these effects are specific to the targeted area and affect behavioral responses [23]. While these studies have resulted in important insights into TMS mechanisms, they were limited to the microscale neurophysiological level and a single brain location. However, it is well understood that TMS effects are widespread, involving several brain networks [24,25]. Furthermore, it is not straightforward to translate the findings from cellular response to EEG. On the other hand, invasive stereo-EEG recordings can measure mesoscale, neurocircuit activity across the brain while preserving excellent spatiotemporal precision. In addition, one can directly translate findings in local field potentials (LFPs) to the macroscale of human EEG.

Here, we record TMS-evoked responses in the NHP brains with implanted depth electrodes along a whole hemisphere. Our experimental setup allows for answering previously unanswered key questions of TMS dose-response and spatial specificity. First, we study the effects of TMS intensity and coil location on TEPs. Second, we measure TMS multisensory co-activation (both auditory and somatosensory effects) through exhaustive control experiments. Our results demonstrate clear TMS-evoked brain responses, that are separable from peripheral effects, with increasing doses corresponding to larger evoked responses. Additionally, we show spatial specificity of these physiological responses. Our findings are crucial for the interpretation of human TMS-EEG studies, rational dose control, and the development of robust markers of target engagement based on TMS-evoked potentials.

## METHODS

### Subjects and surgical procedures

All procedures were approved by the Animal Care and Use Committee of the Nathan Kline Institute. We used two adult non-human primates (*Macaca mulatta*) in this study (Monkey W – female, 5 kg; Monkey H – male, 10 kg). The monkeys were implanted with an MRI compatible PEEK headpost positioned over the occipito-parietal region. Three multi-contact stereo electroencephalography (sEEG) depth electrodes (Ad-tech®, 5 mm spacing) were permanently implanted through a skull incision over the left occipital cortex. Recording electrodes were oriented along the posterior-anterior axis with medial prefrontal cortex, frontal eye field (FEF), auditory (AUD), and temporal cortex (TEM) as the endpoints. In monkey W, an additional electrode targeted the geniculate complex of the thalamus.

### Magnetic resonance imaging (MRI) and electrode localization

For both monkeys, we acquired the T1 and T2 spin echo (SE) sequences using a Siemens® TrioTim 3T scanner with the following parameters: pixel dimensions = 0.5 x 0.5 x 0.5 mm, flip angle = 80°, TR = 2600 ms, TE = 3.55 ms, and TI = 900 ms. Exact electrode positions were identified on a post-implantation MR image, registered to the pre-implantation image, and referenced to the common stereotaxic atlas [26]. See Tables 1-1 and 1-2 for details. MRI registration was done using FSL’s FLIRT package [27–29].

### Transcranial magnetic stimulation (TMS)

We performed the TMS experiments on the two non-human primate subjects in sphinx position. Monkeys were anesthetized with Dexdomitor 0.015mg/kg IM, ketamine 6 mg/kg IM, atropine 0.045 mg/kg IM, followed by 1-2% isoflurane. In all experimental sessions, Biphasic single-pulse TMS was delivered using a MagPro X100 (MagVenture®, Denmark) device with a butterfly coil MC-B35. The coil was positioned at locations designated in Fig 1a above the prefrontal cortex (Monkey W and H: position 1-2), premotor cortex (Monkey H: position 3), temporal areas (Monkey W: position 3, Monkey H: position 4), and auditory cortex (Monkey W: position 4, Monkey H: position 5). Stimulation location contralateral to the implanted sEEG electrodes, above the right temporal cortex (Monkey W: position 5, Monkey H: position 6) was used for both monkeys as a control condition. For frontal stimulation locations (Monkey W: position 1-3 and Monkey H: position 1-4), the coil orientation was maintained parallel to anterior-posterior line. Due to the intrusion of the head post on the lateral regions, for lateral stimulation locations (Monkey W: position 4-5 and Monkey H: position 5-6), the coil was oriented approximately 45^0^ with respect to the anterior-posterior line. We ensured that the connectors and the wires to the electrodes were directed away from the coil and the stimulator, and the amplifier were substantially away from the monkey head during stimulation. For each location, we delivered five different TMS intensities in both NHPs: 10%, 25%, 50%, 70%, and 90% of maximum stimulator output (MSO). In addition, in Monkey H we applied 90% of maximum in power mode, equivalent to ∼125% standard MSO. These intensities are linearly related to the current input (coil dI/dt) as follows: 10% MSO = 13 A/μs, 25% MSO = 37 A/μs, 50% MSO = 76 A/μs, 70% MSO = 107 A/μs, 90% MSO = 142 A/μs and 125% MSO = 201 A/μs. Each stimulation block (condition × location × intensity) consisted of ∼80 TMS pulses given every 3–5 seconds.

**Figure 1:**
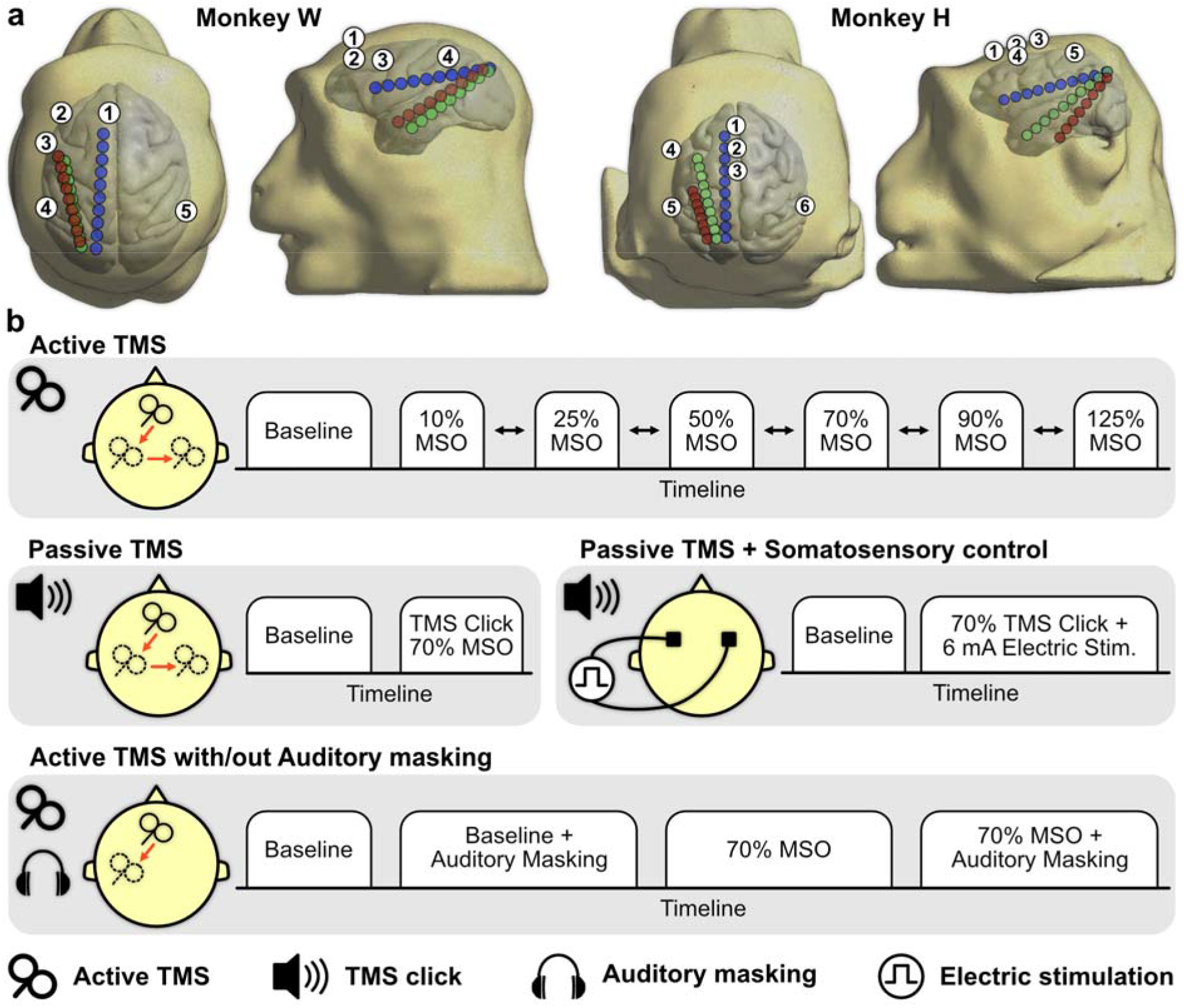
Overview of electrode implantation and experimental setup. a) Contact locations of invasive depth electrodes and TMS coil locations in the reconstructed head and brain surfaces of each monkey. The electrode shafts are directed at frontal eye fields (blue) and auditory (green) and temporal (red; superior temporal in Monkey W and inferior temporal in Monkey H) regions of the brain. TMS is delivered to five locations (numbered) in monkey W and 6 locations in Monkey H. b) Active TMS is delivered at 5 intensities (10, 25, 50, 70, and 90% of the maximum stimulator output, MSO) to both monkeys, with additional power mode setting of 125% MSO in Monkey H. For auditory control, TMS click is delivered at 70% MSO by turning the coil by 90 degrees away from the head, to mimic the auditory response. For somatosensory control, the scalp is stimulated with 6 mA electric pulse while delivering TMS click at 70% MSO. This control was performed at location 2 in monkey H. To control the effects of auditory co-activation, active TMS was delivered at 70% MSO with and without auditory masking at locations 1 and 5 in Monkey H.

We implemented the following experimental conditions (Figure 1b): I) Active TMS, II) Passive TMS (produces auditory click associated with TMS), III) Passive TMS with somatosensory control (adds focal electric pulse to the scalp), and IV) Active TMS with auditory masking (continuous auditory white noise masking auditory click associated with TMS).

In the passive TMS condition, we turned the TMS coil away from the head to solely mimic the TMS auditory response due to the click sound of the coil without direct brain stimulation. For auditory/somatosensory co-stimulation, we used MagVenture Cool-B65 A/P coil that concurrently provides electric current stimulation to the scalp. To study whether the current stimulation alone elicits peripheral effects, we used the sham coil in combination with the current stimulation over stimulation location 1. We delivered auditory click at 70% MSO along with 6 mA electrical stimulation on the scalp using pairs of 3M disc electrodes of 10mm diameter (3M®, Saint Paul, MN, USA) placed across the region of interest. This sham stimulation approach follows Siebner et al. [17]. Given that the cutaneous perception of electrical stimulation due to active TMS and somatosensory control cannot be assessed subjectively through feedback from nonhuman primates, we visually assessed and confirmed equivalent scalp and shoulder twitches during active and sham conditions.

For auditory masking, we created a masking acoustic noise using the TMS Adaptable Auditory Control tool [30]. The sound pressure level of the masking noise was set to 80 dB for the above stimulation intensities. Sound pressure level was measured at the ear canal using a sound meter (Bruel and Kjaer®, Nærum, Denmark). The masking noise was presented via two speakers placed 50 mm from the head on each side. The masking was applied concurrently with active TMS at stimulation locations 1 and 5 with the intensity of 70% MSO.

### Local field potentials (LFPs)

We recorded the local field potentials (LFPs) using 32-channel ActiveTwo amplifier (Cortech®, Wilmington, NC, USA) at a sampling rate of 40 kHz. For both monkeys, contacts in the occipital cortex (proximal to the headpost) were chosen as the reference and the ground (9^th^ and 11^th^ contacts from the front in the FEF electrode respectively). To prevent the electric current induction in the electrode contacts due to the magnetic field, we made sure that the recording setup, including the electrode contacts and the electrical cables, were free of loops. Furthermore, high input resistance of the amplifier (1 GΩ) would minimize induced currents to negligible levels, even if they occur.

Data preprocessing was done in MATLAB (MathWorks® Inc., Natick, MA, USA) using the FieldTrip toolbox [31]. The data acquired from the amplifier were preprocessed by separating data into epochs time-locked to TMS delivery (t = 0). The epoch length was 4 s (2 s before and 2 s after TMS delivery). The epochs were then detrended and demeaned. In the next step, we removed the TMS pulse artifact (−0.5 ms to 0.4 ms) and the TMS-induced muscle artifact (0.4 ms to 25 ms in both monkeys) by padding the time windows. The time windows to be excluded were determined by the visual inspection of the LFPs and concurrent electromyographic activity recorded from scalp (see Figure 1-1). We interpolated the excluded time window using cubic Hermite interpolating polynomial (pchip) with pre and post data segments of 200 ms. We filtered the resulting LFPs using a bandstop Butterworth filter of order 2 with cutoff frequencies at 57 and 63 Hz. Noisy contacts and trials were excluded through visual inspection. 11 ± 6 (mean ± SD) trials were excluded per experimental block. We performed independent component analysis to manually remove residual artifacts (one component on average per experimental block, maximum of 4). The resulting time series data was then resampled to 1 kHz and bandpass filtered from 1–50 Hz using a 4th order Butterworth filter, followed by baseline correction. See Figure 1-1. We processed baseline data by splitting the traces into 4 s epochs with 50% overlap. We followed the same preprocessing steps for baseline data except for artifact removal and interpolation. For subsequent analysis, we excluded the sEEG contacts located in white matter or outside the brain.

### Electric field modeling

We modeled the electric fields induced by TMS using SimNIBS 3 [32]. We extracted the gray matter and white matter surfaces using a modified Human Connectome Project pipeline for non-human primates [33]. The surfaces for scalp, skull, and cerebrospinal fluid were created by manual segmentation. Once the complete head mesh was created, we ran simulations for the TMS coil positions used experimentally. We assigned fixed conductivities to each layer: 0.126 S/m (white matter), 0.275 S/m (gray matter), 1.654 S/m (cerebrospinal fluid), 0.01 S/m (skull), 0.465 S/m (scalp) [34]. The electric fields were calculated in a quasi-static regime [35]. Through electric field simulations, we sought to visualize the affected brain region and the extent of activation. We computed the robust maximum (99.95^th^ percentile) of the electric field magnitude to remove numerical inaccuracies. To calculate the spatial similarity of electric field distributions on gray matter surface, we first obtained the binary mask by thresholding the electric field magnitude at 50% of the robust maximum. Then, we computed the Sørensen-Dice similarity coefficients on pairs of gray matter surfaces.

### Data Analysis

We performed a three-step analysis on the electrophysiological data. First, we analyzed the dose dependency of identified TEPs. Second, we investigated the effect of coil-to-contact distance on the TEP response. Finally, we evaluated the region-specific differences in LFPs and TEP components in the presence of multisensory co-activation.

Following the data preprocessing, we estimated the significance of time-locked LFPs or TMS-evoked potentials (TEPs). The average time-locked activity was calculated for each experimental block. For each stimulation location, we performed a non-parametric cluster-based permutation test [31] (p < 0.05) to investigate whether a significant main effect of stimulation intensity is present following TMS (25-400 ms after stimulation pulse) relative to the baseline. The F statistic was calculated for each contact/time pair and clustered according to their temporal adjacency. We used the sum of the F values within a cluster as a cluster-level statistic and compared it against a permutation distribution of 1000 permutations to determine the p value. The highest F value per cluster is reported in Results. The contacts in which the effect was present during at least one stimulation intensity were considered to be ‘responsive’ contacts.

Identified significant TEPs were named according to the timepoint of the maximum deflection (e.g., TEP with the maximum negative deflection at 50 ms is N50). We extracted the N50 component as the minimum amplitude of the LFP in the time window of 40-70 ms from TMS onset, per each trial. To analyze the dose dependency of the TEP components, we extracted N50 for all intensities, from all responsive contacts, and from each stimulation location. TEP amplitudes were normalized to the maximum at each stimulation location. The responses from both monkeys, responsive contacts and stimulation locations were pooled and fit to a four-parameter logistic function,

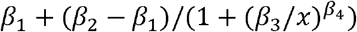

with stimulation intensity as the independent variable and the TEP response as the dependent variable.

Next, we studied the effect of stimulation location on the TEP response through a distance-response relationship. We calculated the Euclidian distances from the TMS coil center projected on the scalp to each electrode contact at each stimulation location. Here, we excluded the contacts without TEP response relative to the baseline (N50 amplitude is within the 99^th^ percentile of the mean LFP amplitude). We assessed the relationship between the distance and response by fitting a linear (y = a*x + b) and an exponential decay function (y = a + b/x^3^). The Goodness-of-Fit was estimated using adjusted R^2^.

Furthermore, we studied the LFPs for the different control conditions. To investigate the peripheral auditory effects, we compared active TMS with and without auditory masking. First, we calculated the averaged total response by combining the trials from all responsive contacts in both tested stimulation locations (locations 1 and 5). To narrow down the auditory and non-auditory responses, we grouped the response of auditory and non-auditory contacts separately and calculated the average response for each condition i.e., masked, and non-masked.

To test whether there’s a statistically significant effect of masking on the TEP responses, we extracted the TEP components from all the trials, and performed Wilcoxon rank sum test (p < 0.05) on the total, auditory, and non-auditory TEP responses.

Finally, we studied the effects of auditory click and peripheral somatosensory effects by comparing active TMS, passive TMS and passive TMS + electrical stimulation. Similar to the auditory masking analysis, we calculated the total averaged response from all responsive contacts. Then we grouped the auditory and non-auditory contacts and calculated the average response. To study the effect of auditory click and electrical stimulation on TEP responses, we extracted TEP responses from all contacts and performed Kruskal-Wallis test (p < 0.05) on total, auditory and non-auditory TEP responses. We performed follow-up multiple comparisons to evaluate the significant differences across the conditions.

## RESULTS

### TMS induced neural effects show a dose dependency

To study the stimulation intensity (dose) dependent effects of TMS on time-locked local field potentials (LFPs), we first identified time windows that showed a significantly different LFP amplitude after stimulation pulse relative to the baseline for at least one stimulation intensity. This was done at five and six separate stimulation locations for two monkeys (named W and H), respectively). The responses were tested using nonparametric cluster-based permutation test. The TMS-evoked LFPs from both monkeys show a high amplitude activity at ∼25-100 ms after stimulation pulse followed by a rebound. In both monkeys, this pattern of activity is conspicuous at the stronger stimulation intensities (≥ 50% of maximum stimulator output, MSO).

In monkey W, upon stimulation of location 1, three frontal contacts showed significant responses (29-102 ms, F_max_ = 7.93-11.89, p = 0.003-0.037). Stimulation of location 2 showed significance in two frontal contacts (25-99 ms, F_max_ = 6.45-9.53, p = 0.005-0.046). Stimulation of location 3 and 4 resulted in significant responses across four (25-99 ms, F_max_ = 12.36-24.16, p = 9.9×10^−4^ - 0.0017) and five (25-133 ms, F_max_ = 13.27-49.52, p = 9.9×10^−4^ - 0.003) frontal contacts respectively. The contacts mentioned above were located in the corpus striatum and anterior cingulate cortex (see Table 1-1 for detail). Stimulation location 4 also led to significant responses in the two contacts within the auditory belt (25-91 ms, F_max_ = 19.93-23.92, p = 9.9×10^−4^ - 0.004) and a cluster of four contacts within the superior temporal plane (25-111 ms, F_max_ = 9.35-21.65, p = 9.9×10^−4^ - 0.014). See Figure 2-1 and Table 2-1. In monkey H, the main stimulation locations 1, 2, 4, and 5 showed early latency responses in the contacts that are in the orbitofrontal cortex, corpus striatum, and anterior cingulate. i.e., three contacts in location 1 (25-93 ms, F_max_ = 6.63-12.95, p = 0.0015-0.05), three contacts in location 2 (27-93 ms, F_max_ = 7.39-23.31, p = 9.9×10^−4^-0.03), three contacts in location 4 (25-130 ms, F_max_ = 8.08-18.25, p = 9.9×10^−4^-0.042) and five contacts in location 5 (25-133 ms, F_max_ = 12.17-44.03, p = 9.9×10^−4^). For all main stimulation locations, the contacts in the auditory cortex saw early latency responses (25-145 ms, F_max_ = 6.64-22.61, p = 9.9×10^−4^-0.045). Locations 1, 3, 4, and 5 also showed significant responses in the contacts in the ventral insular region (25-161 ms, F_max_ = 4.40-18.85, p = 9.9×10^−4^-0.045). Furthermore, all main stimulation locations resulted in significant responses in the superior temporal and the middle temporal regions while the responses due to stimulating location 5 resulted in longer time windows (Locations 1, 2, 3, and 4: 25-117 ms, F_max_ = 5.46-11.79, p = 0.003-0.041 and location 5: 33-174 ms, F_max_ = 10.03-31.64, p = 9.9×10^−4^). See Figure 2-1 and Table 2-2. The control stimulation location (location 5) in monkey W did not elicit a significant response in the LFPs. However, in monkey H, control stimulation location (location 6) resulted in significant early latency responses in the frontal contacts. The LFPs resulting from stimulating the control location in comparison to its ipsilateral counterpart is shown in Figure 2-2.

**Figure 2:**
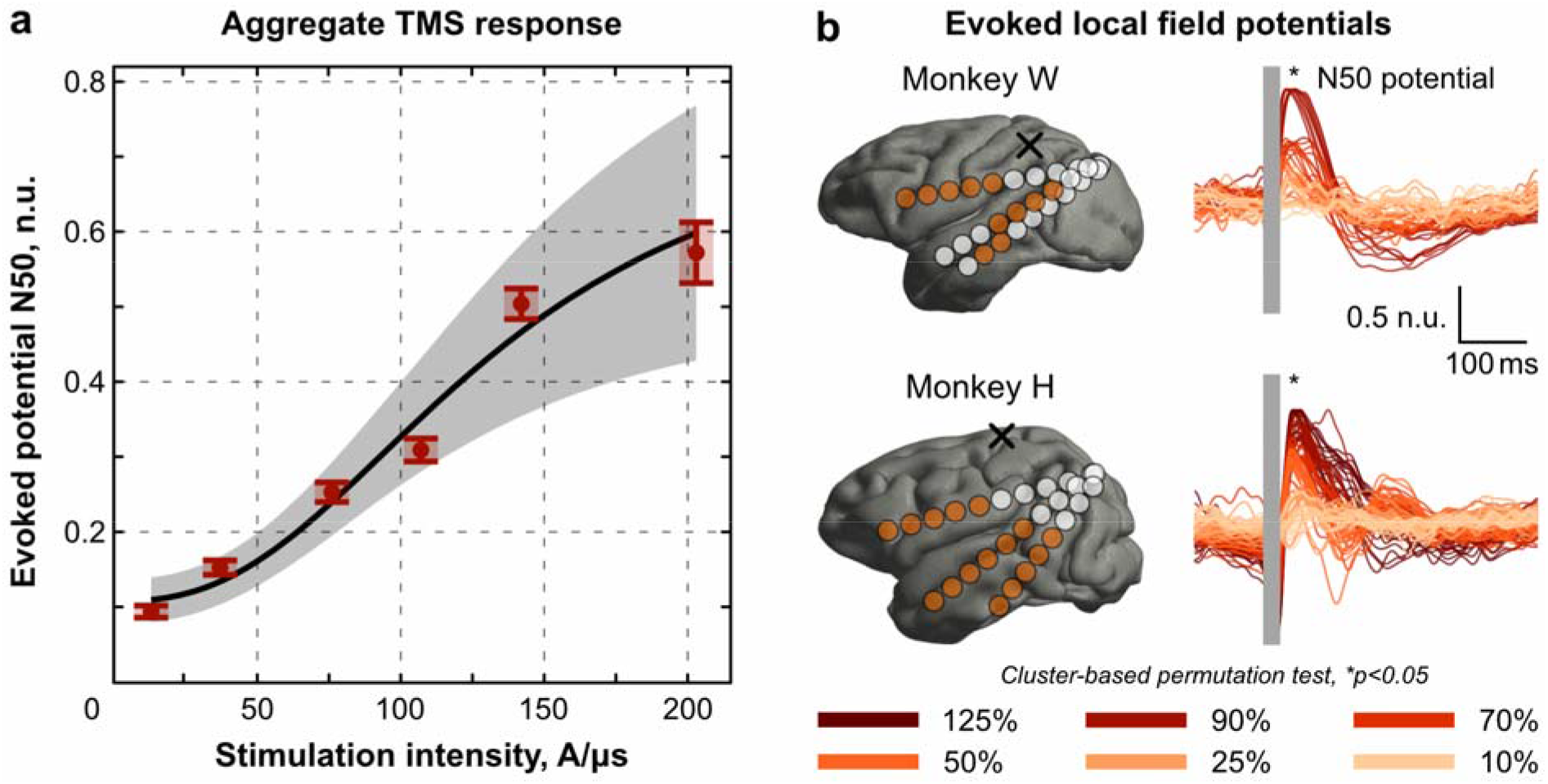
TMS-induced early evoked potential N50. a) Aggregated magnitude of early evoked potential N50 in all responsive contacts (significant for at least one stimulation intensity) in both monkeys. The N50 amplitude is normalized per stimulation location per monkey. The curve shows the sigmoidal (logistic) dose-response function with the upper and lower confidence bounds (shaded). The mean squared error of the fit is 0.018. b) Left. The 3D rendering of the brain with the recording contacts (responsive contacts are in dark orange) in both monkeys for a representative stimulation location. Right. normalized local field potentials (overlayed responsive contacts). Responsive contacts were determined by nonparametric cluster-based permutation test, *p < 0.05.

In the early latency time windows, the LFPs showed a maximum deflection ∼50 ms after TMS delivery. This N50 component demonstrated a pronounced increase in its magnitude with increasing stimulation intensity in the lateral-most stimulation location (locations 3 and 4 in monkey W and locations 4 and 5 in monkey H) in both monkeys. In locations 1 and 2 in monkey W and locations 1, 2 and 3 in monkey H, the N50 component showed a robust response for higher stimulation intensities (90% MSO and 125% MSO). We extracted the N50 component from all responsive contacts across all three electrodes from both monkeys to investigate the dose dependent behavior. The number of total responsive contacts was 20 (3+2+4+11) and 57 (11+10+8+12+16) out of 108 and 135 in monkeys W and H, correspondingly. The N50 components were normalized to the maximum N50 value for each stimulation location. To estimate the overall dose dependency of N50, we fit a four-parameter sigmoid (logistic) curve with TMS dose as the independent variable and the normalized N50 deflection as the dependent variable (Figure 2a). The mean squared error of the fit was 0.018.

### Effect of coil location and response-distance relationship

We performed computational simulations to evaluate the spatial extent of the TMS-induced electric field in the brain under different coil positions (Figure 3a). As shown in Figure 3b, in monkey W, pairwise Sørensen-Dice similarity coefficient showed anticipated moderate overlap between the main stimulation locations 2-4 (DICE score: 0.19-0.56). In monkey H, the fronto-medial stimulation locations 1, 2, and 3 showed high spatial similarities (DICE score: 0.54-0.75), while the lateral locations 4 and 5 showed moderate similarity between themselves (DICE score: 0.36). The control stimulation location in monkey W showed no impact on the recorded hemisphere (DICE scores with the main stimulation locations < 0.01). In monkey H, the control stimulation location showed little-to-no impact on recorded hemisphere (DICE scores with the stimulation locations 1, 2, 3, and 4 was < 0.15) except for stimulation location 3 (DICE score of 0.29).

**Figure 3:**
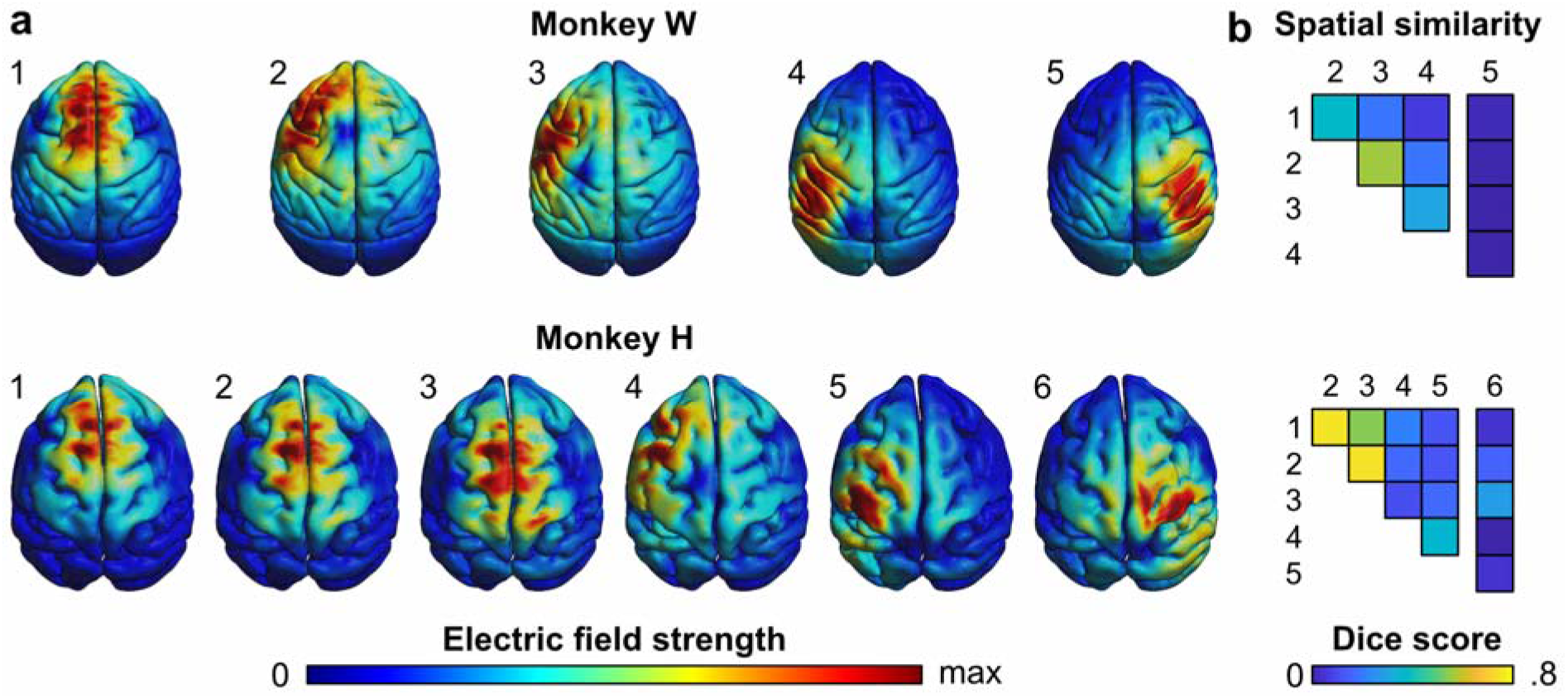
TMS-induced electric fields on cortical surfaces. a) The simulations of the TMS-induced electric field strength on the brain surface are shown in Monkeys W and H for each coil location. The electric field strength is color-coded in relative units with respect to the maximum value. In Monkey W, locations 1 (anterior medial) and 2-4 (lateral) constitute main experimental conditions and location 5 is the contralateral control condition. In Monkey H, locations 1-3 (medial) and 4-5 (lateral) are main experimental conditions, and contralateral location 6 is the control. b) The spatial similarity between TMS-induced electric fields as quantified by Sørensen-Dice similarity coefficient. In monkey W (top panel), main stimulation locations demonstrated moderate spatial similarity between pairs (DICE score: 0.19-0.56) while the control stimulation location showed no impact on the recorded hemisphere. In monkey H (bottom panel), the stimulation locations 1, 2, and 3 showed high spatial similarities (DICE score: 0.54 – 0.75), while the lateral locations 4 and 5 showed moderate similarity between themselves (DICE score: 0.36). The control stimulation location showed little impact on the recorded hemisphere except for stimulation location 3 (DICE score: 0.29).

We next explored the relationship between stimulation location and the N50 response. We focus on the high intensity (90% MSO) condition as it elicits a robust response across most contacts for both monkeys. We combined the stimulation locations 1, 2, and 3 in monkey W and locations 1, 2, 3, and 4 in monkey H for this analysis that were comparable in coil orientation. The lateral most locations (4 in monkey W and 5 in monkey H) were excluded given the different coil orientation. Thus, N = 33 (15+7+11) and N = 49 (14+11+11+13) contacts in both monkeys from the said stimulation locations were included in this analysis. To assess the relationship between the N50 response and the coil-to-contact distance, we fit linear and exponential decay models (Figure 4 and 4-1. A negative linear relationship between the N50 response and the coil-to-contact distance in monkeys W (R^2^-adjusted = 0.51) and H (R^2^-adjusted = 0.23) is significant but less accurate than a decaying relationship. The exponential decay fit is much better for monkey W (R^2^-adjusted = 0.74) and similar to the linear fit in monkey H (R^2^-adjusted = 0.18).

**Figure 4:**
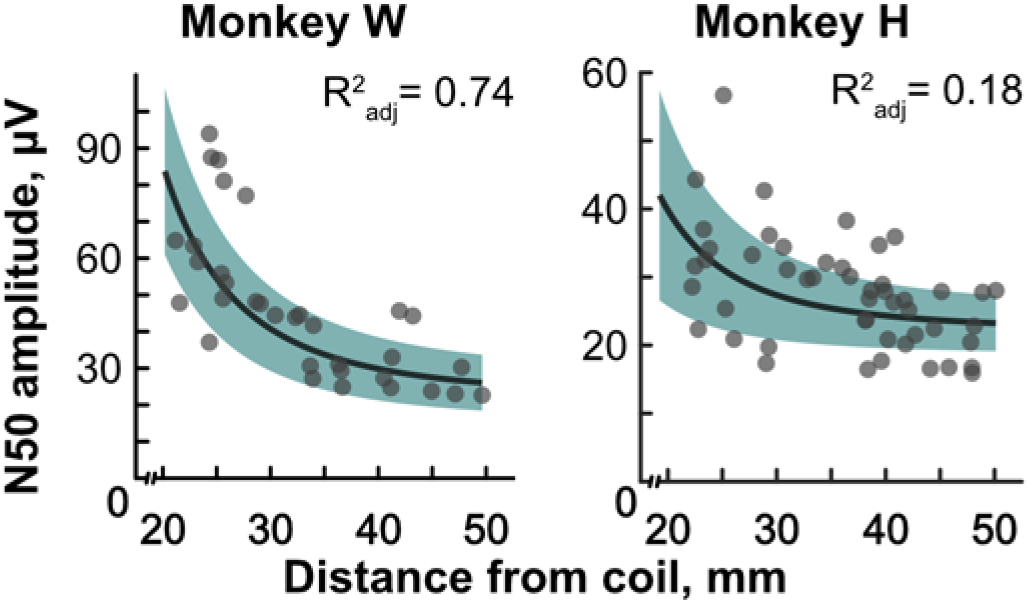
Relationship between N50 response and the coil-to-contact distance. Scatter plots show the amplitude of the N50 response against the coil-to-contact distance for each stimulation location. Stimulation locations 1-3 from monkey W and 1-4 from monkey H are included in this analysis. The relationship between the N50 amplitude and the coil-to-contact distance in monkeys W (R2-adjusted = 0.74) and H (R2-adjusted = 0.18) show an exponential decay.

### TMS induces direct and peripheral stimulation effects in the auditory cortex

We performed three control experiments to study the effects of peripheral auditory and somatosensory co-stimulation. First, we compared auditory and non-auditory regions during active TMS and active TMS with a masking noise, which suppresses the TMS-associated click sound. For this control, we stimulated locations 1 and 5 in monkey H and extracted LFPs from all contacts that were responsive for both locations. We computed the average LFP responses across all contacts and separately for contacts in auditory and non-auditory brain areas for masked and unmasked conditions separately. The average non-auditory response did not show a visible change between the masked and unmasked conditions. However, the average response and average auditory response show a reduction in amplitude in the early time window in the masked condition compared to the unmasked condition (Fig 5a). We further investigated this by extracting the N50 components from the LFPs. The components were normalized to the maximum at each stimulation location. We performed Wilcoxon rank sum test to compare the N50 responses between masked and unmasked conditions for both regions. We found that non-auditory regions are immune to masking (Z = 1.09, p = 0.27) whereas auditory regions show a significantly lower N50 response during masking (Z = 5.36, p = 8.1 × 10^−8^). The average response also shows a decreased response when masked (Z = 3.47, p = 5.1 × 10^−4^). The results suggest that TMS induces direct effects on neuronal populations in the stimulation area due to electric field, while also inducing separate response in the auditory cortex due to stimulation click sound.

**Figure 5:**
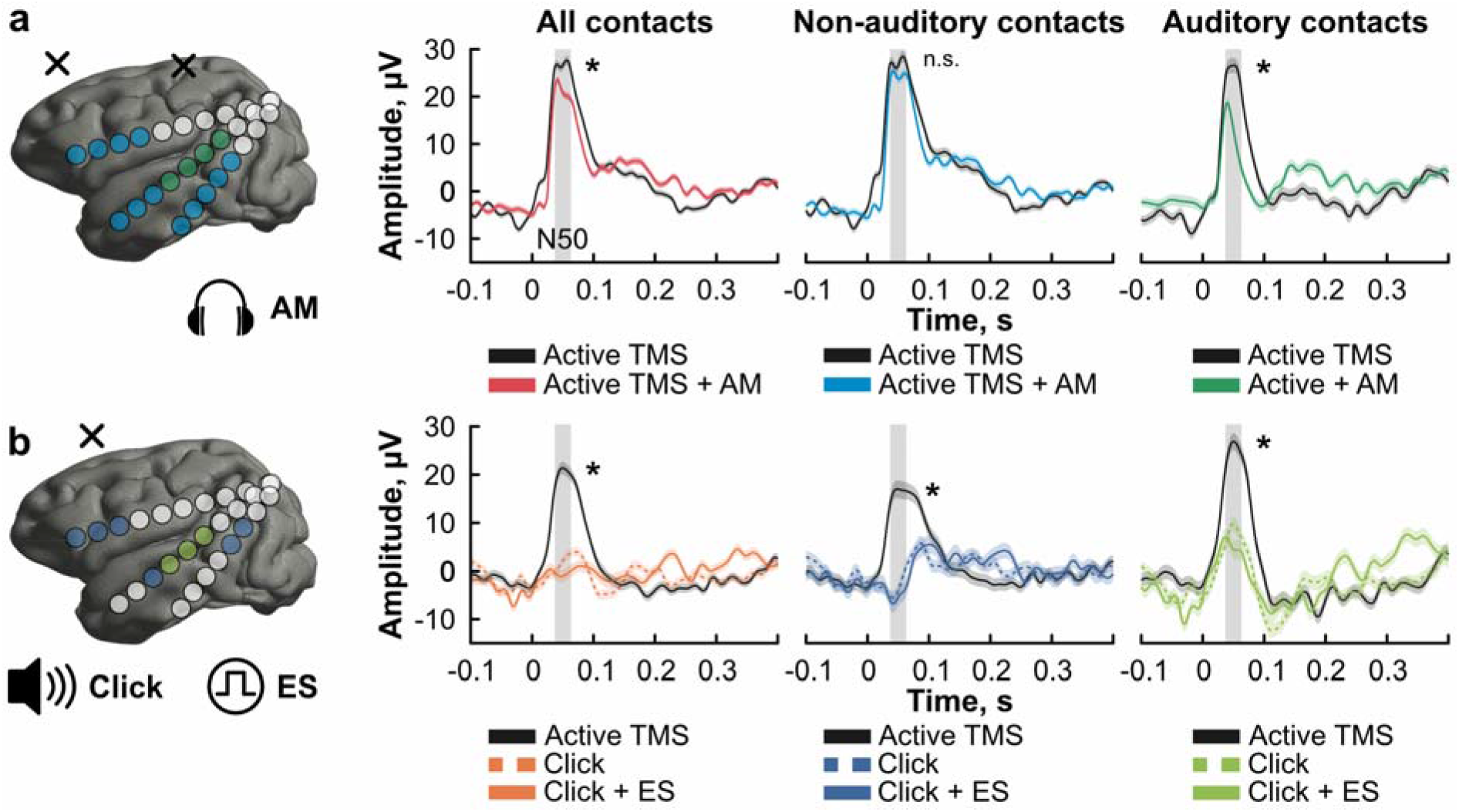
TMS-induced direct neural effects are separable from peripheral auditory and somatosensory activation. Total average response, average non-auditory response, and average auditory response are calculated from the responsive contacts. The responsive contacts are shaded in blue (non-auditory) and green (auditory). The TMS coil locations are indicated by X above the cortical surface. The N50 responses are extracted on a trial basis for comparison across controls. a) Auditory masking results in diminished N50 response localized to the auditory region when active TMS is delivered. Wilcoxon rank sum test indicates significantly lower N50 response in the average response (p = 5.1 x 10^−4^). When the responses are clustered into auditory and non-auditory regions, the same test indicates that the N50 response in the auditory region shows a significantly lower response (p = 8.1 x 10^−8^) in the masked condition compared to unmasked when delivering active TMS. The test returns no significant difference for the N50 response between masked and unmasked conditions in the non-auditory regions. b) Auditory click and auditory click + electrical stimulation do not induce N50 responses comparable to active TMS. Kruskal-Wallis tests for N50 component for total response, non-auditory response and auditory response showed a significant difference across the three conditions (Total: χ^2^ = 114.68, p = 1.25 x 10^−25^, Auditory: χ^2^ = 30.56, p = 2.31 x 10^−7^, non-auditory: χ^2^ = 89.15, p = 4.38 x 10^−20^). Follow-up multiple comparisons tests revealed that the total N50 response was significantly different for the active TMS vs. click (p = 9.56 x 10^−9^) and active TMS vs. click with electric stimulation (ES; p = 9.56 x 10^−9^). The auditory N50 response was also significantly different for the active TMS vs. click (p = 5.29 x 10^−5^) and active TMS vs. click with ES (p = 7.26 x 10^−7^). A similar observation was made on non-auditory N50 response where the active TMS vs. click (p = 9.74 x 10^−10^) and active TMS vs. click with ES (p = 9.56 x 10^−10^) were significantly different. n.s. = not significant, *p < 0.05.

### Auditory controls show a localized response in the auditory cortex

We compared the effects of auditory click control condition and auditory click with electric scalp stimulation to active TMS (Figure 5b). As in the previous section, we considered the total, auditory, and non-auditory responses across conditions. The average LFPs from all three groups showed a pronounced N50 response in the active TMS condition (Figure 5b). Click only and click with electric stimulation did not show a pronounced N50 response in the total and nonauditory groups. However, both showed a moderate N50 response in the auditory group that was less than that of the active TMS condition. We performed Kruskal-Wallis test on the extracted N50 responses to test for a significant main effects of stimulation condition (active TMS, click only, and click + electric stimulation). For all three groups, we observed significance (Total: X^2^= 114.68, p = 1.25 × 10^−25^; non-auditory: X^2^ = 89.15, p = 4.38 × 10^−20^; auditory: X^2^ = 30.56, p = 2.31 × 10^−7^) for at least one stimulation condition. Follow-up multiple comparisons tests indicated that active TMS evokes a significantly higher N50 response than both control conditions across all groups. Comparison of total response showed significance for the active TMS vs. click (p = 9.56 × 10^−9^) and active TMS vs. click with electric stimulation (p = 9.56 × 10^−9^).

Similarly, response in the auditory cluster showed significance for the active TMS vs. click (p = 5.29 × 10^−5^) and active TMS vs. click with electric stimulation (p = 7.26 × 10^−7^). Same pairs of conditions were significant in the non-auditory cluster: active TMS vs. click (p = 9.74 × 10^−10^) and active TMS vs. click with electric stimulation (p = 9.56 × 10^−10^). This result confirms that the sham protocol used to mimic auditory click and scalp stimulation does not induce TMS effects in non-auditory regions. Furthermore, scalp stimulation did not induce any significant effect on total or non-auditory response, confirming that somatosensory co-activation did not affect the recorded brain regions. In the auditory regions, the auditory click results in N50 response for both control conditions, which is significantly less than in the active TMS condition. This corroborates the observation made in the auditory masking condition that effects on auditory regions include TMS-induced direct effects as well as peripherally induced effects due to auditory activation.

## DISCUSSION

We investigated how single-pulse transcranial magnetic stimulation (TMS) affects mesoscale level neuronal activity in non-human primates depending on TMS intensity and coil position. Further, we employed several control conditions to dissociate the target effects in the brain from peripheral auditory and somatosensory side effects. We found that early evoked potentials, occurring ∼50 ms after TMS pulse, characterize a direct local neural response to stimulation. These early local field potentials demonstrate a characteristic, sigmoidal dose-response dependency and inversely relate to the distance from the brain area to the stimulation coil. This direct brain response is well dissociable from acoustic co-stimulation, particularly evident in the auditory cortex, and somatosensory co-stimulation, which doesn’t induce specific early evoked components. These findings provide the basis for concurrent TMS-EEG studies in humans and put forward early evoked potentials as a clinically-relevant target engagement biomarker.

### TMS-evoked potentials and dose-response

We found that TMS stimulation results in TEPs characterized by high amplitude and short latency. In both monkeys, the early activity expressed itself as a positive peak at ∼50 ms. In monkey W, we saw a later rebound component at the highest stimulation intensity. Comparable observation has previously been reported in TMS-EEG studies in humans in unconscious or anesthetized states [12,36,37]. During such states, the TEPs are localized to the stimulation region, which results in less complex brain activity. However, in attentional states, the long-range brain connectivity results in complex TEPs with multiple peaks. The early response we observed here was robust in several brain areas across multiple TMS coil locations. While fronto-medial stimulation locations are only effective at high doses (e.g., 90% MSO), lateral stimulation locations showed a characteristic, sigmoidal dose-response dependency with increasing TMS intensity. In particular, we found that early LFP component (N50) captures a mesoscale brain response not reported in the nonhuman primate literature so far. Nevertheless, the effects of single pulse TMS on the visual cortex of anesthetized cats have been studied with electrophysiological recordings and optical imaging [38,39]. While these studies provide important insights into complex TEPs patterns, research in nonhuman primates enables straightforward translation to humans [21]. Two groups have studied the effects of TMS on the microscale neuronal level in nonhuman primates. One group examined single unit activity during stimulation in different types of neurons [22]. Example neurons showed bursts of activity at short latencies (< 100 ms), especially around 50 ms. The other group showed that the majority of single units demonstrated increased short latency (10 – 40 ms after TMS delivery) spiking activity in parietal neurons during task-related activity, while some neurons demonstrated excitation-inhibition-excitation pattern [23]. While the summation of single neural firing should result in the population-level response, our study directly shows the aggregate LFP activity in neural circuits and its potentiation approximately 50 ms after TMS delivery. Hence the N50 component captures direct physiological effect of TMS.

To ensure that the TEPs are not a consequence of currents induced by inductive loops in the recording setup, we ensured that the contacts of the electrodes and the electrical cables were free of loops. Furthermore, the high input impedance of the amplifier results in negligible currents compared to the biological currents that gives ride to the LFPs. An interesting question is whether the TEPs observed here are due to direct neural stimulation close to the recording sites, propagated from more superficial brain areas via neural circuits, or both. The TMS-induced electric fields generate maximum at the neocortex; however, a substantial part of the electric field in the small NHP brain also reaches deep areas. The cross sections of the electric field magnitudes at each electrode for each stimulation location are shown in Figure 3-1. Thus, our data cannot directly distinguish between the abovementioned possibilities, which remain an exciting avenue for future research.

We also observed that the lateral-most stimulation locations in both monkeys (location 4 in W and location 5 in H) showed activation in the distal contacts compared to the proximal ones. We speculate that this could be due to a couple of reasons: 1) The coil orientation used for these two locations were different from the rest due to the headpost placement, which could have resulted in activation of deeper, frontal regions and 2) The lack of activation in the proximal contacts could be due to the placement of some of these contacts in the WM (or WM and GM interface – refer Tables 1-1 and 1-2). We did not observe the same LFPs in these contacts and they were excluded from the analysis.

Dose selection is a vital ingredient in TMS application, whether in research or therapy. Rational dose selection enables achieving desired treatment outcomes [40], and eliciting desired activity in the brain when probing brain networks. Coupling the stimulation dose with a biomarker that reflects direct TMS-induced neural activity is a key prerequisite for developing optimized dose control. In human TMS studies targeting the motor cortex, the effect of TMS stimulation intensity has mostly been studied using motor-evoked potentials (MEPs) [41]. Given that MEPs are an indirect readout of cortical reactivity in the form of a muscle response, the effects of TMS intensity can be quantified easily. However, in brain regions pertaining to cognitive processes, for example, prefrontal or parietal cortex, such dose-response relationships have not been established with easily accessible readouts. In such cases, TEPs can serve as important biomarkers that explain the effects of stimulation intensity and connect them with behavioral responses. In previous human work, the intensity dependency of TEPs in human PFC was demonstrated using TMS-EEG [42]. However, this study was limited to a single brain region and did not disentangle direct TMS effects from multisensory co-activation.

### Spatial dependency

We used electric field simulations to study the effect of coil position on the field distribution. The Sørensen-Dice similarity coefficient analysis indicated similarity in electric field distributions between stimulation locations that are on the hemisphere ipsilateral to recording electrodes. The smaller size of the monkey brain and the range of coil movement that was ≤ 3 cm between adjacent stimulation locations, lead to overlapping field distributions. Control location 5 in monkey H scarcely overlapped with distributions from other locations. However, in monkey H, two medial stimulation locations had overlapping distributions with the control location. Furthermore, the electric field distribution when stimulation location 6 in monkey head also showed electric fields reaching the medial electrode in the left hemisphere. This can explain the response we observed in LFPs when stimulating location 6. As the monkey brain is considerably smaller than the human brain, the percentage of the brain volume activated by TMS is higher in monkeys than in humans [21]. Thus, the spatial specificity of TMS is likely reduced to what would be expected in humans.

Investigating the spatial dependency of TEP responses, we observed a relationship between the N50 amplitude and the distance from the TMS coil to the recording contact. An inverse linear model and an exponential decay model can explain the seen relationship. Yet, the decay model captures this effect better, especially in monkey W. This observation agrees with the nature of the magnetic field, which is rapidly decaying in magnitude with increasing distance from its source.

The effect of coil-to-cortex distance on TMS effects on DLPFC [43] and motor cortex [44,45] has been studied previously across subjects. While Romero et al. [23] showed that TMS elicits spiking activity with a very high spatial specificity, we observed more widespread neural effects on the brain. The key differences in this study and our study potentially led to different observations: I) Their stimulation paradigm is limited to one brain region, hence is limited in explaining effects arising from stimulating other regions, II) they record from a single brain region and therefore cannot capture the induced effects that are propagated to connected regions, and III) the difference of the scale of recording, i.e., single units vs neural circuits.

### Multisensory co-activation

We incorporated several control conditions to disentangle direct neural effects from peripheral sensory effects. We conclude that auditory masking only affects the response in auditory regions confirming that a fraction of the response due to TMS is in fact auditory evoked in nature. Similarly, the difference of responses in active TMS and passive TMS (auditory click) conditions confirms the above observation. We did not observe a visible response in nonauditory regions during passive TMS. Electrical stimulation of the scalp (coupled with passive TMS) to control for somatosensory activation did not produce a significant response in the nonauditory regions. Since the depth electrodes did not have contacts in the somatosensory cortex, we could not determine whether a response localized to that region was present during electrical stimulation. The evoked potentials we observed during passive TMS and electrical stimulation did not elicit long latency responses. In fact, the somatosensory evoked potentials in anesthetized monkey preparations have shown to be short latency in nature and lasting up to 25 ms [46]. Therefore, it should not impact our findings in the responses appearing after 25 ms. Research of different anesthetics in humans shows the normal modulation of middle latency auditory evoked potentials at 10-100 ms after stimulus delivery [47]. These modulatory effects are also shown for somatosensory sensation [48]. Hence, our analysis is confined to the TEP components that show modulatory effects for the given conditions. Present results have implications in TMS-EEG studies. Interpreting TEP components and assessing their origins is challenging due to limitations of EEG recordings [17]. A recent study has attempted to disentangle peripheral activations from direct TMS effects using similar controls [20]. The study has compared the TEPs across conditions in the vicinity of a single stimulation site. Despite similar observations being made on modulation early evoked activity, direct recordings from auditory regions in our study give us valuable insights about direct and peripheral TMS effects on auditory and nonauditory regions.

### Non-human primate – human translation

The main TEP component we observed in this study can be compared with some of the human TMS-EEG studies. Several human TMS-EEG studies have reported N100 response resulting from stimulation in the vicinity of M1 [49,50]. The N100 component has also been observed in TEPs resulting from prefrontal cortex stimulation [13,42]. This component has been implicated as a neurophysiological marker of cortical inhibition through MEP measurements [49,51]. The disparity in latency could be attributed to the differences in neural dynamics, gross functional anatomy and cellular physiology between humans and monkeys [52]. However, there is also evidence for TMS evoked responses in humans that are commonly characterized by early potentials such as cortical N40 and occur on the same timescale as we observed here [53,54]. Importantly, the distinction between recording modalities also contributes to differences in latency. Furthermore, using anesthetized preparations of non-human primates also contributes to such differences. In human literature, several TMS-EEG studies reveal complex TEP waveforms during attentional states. These include peaks at 15, 30, 45, 60, and 100 ms during motor cortex stimulation [18,55] and peaks at 27, 40, 60,100, and 180 ms following prefrontal stimulation [56,57]. However, in sleep or unconscious states, the TEP components are less complex, more localized, and higher in amplitude [12,36,37]. We observe response patterns in two anesthetized monkeys equivalent to the sleep or unconscious findings in humans. Furthermore, the human data on subcortical ERPs is very scarce and it is not clear if the subcortical ERP we observed here is more or less complex than its human counterpart. While TMS induced effects are state dependent, anesthetized models still provide important mechanistic understanding about neural effects induced by TMS. Previous studies have used anesthetized monkey models to explain the activity of visual cortex when presented with visual stimuli [58] and effects of repetitive TMS using functional imaging [59,60]. Nevertheless, future work is needed to establish N50 as a biomarker in humans and the present work lays significant advances towards this goal.

### Conclusion

Understanding the relationship between TMS dose and physiological responses is crucial for optimizing TMS parameters to achieve desired clinical outcomes, elicit network activity, and induce plasticity. Furthermore, understanding the origins of evoked components in non-invasive EEG and the correct interpretation of TEPs in the presence of multisensory coactivation improve their utility as biomarkers for target engagement. Our study provides important findings regarding the dose-dependent physiological effects of TMS, the role of stimulation location and the distance between the target brain region and the coil position, and the impact of multisensory coactivation due to TMS on evoked potentials. Together, present results provide a mechanistic basis for human TMS-EEG studies and TMS biomarkers for clinical therapy.

## Acknowledgements

Research presented here was supported by the National Institutes of Health grants R01NS109498, RF1MH117428, and K99MH128454, and the University of Minnesota MnDRIVE Initiative.

## Supplementary information

Supplementary material is available for this paper online.

