## Supplemental figures and tables for "Dissociation of direct and peripheral transcranial magnetic stimulation effects in nonhuman primates"

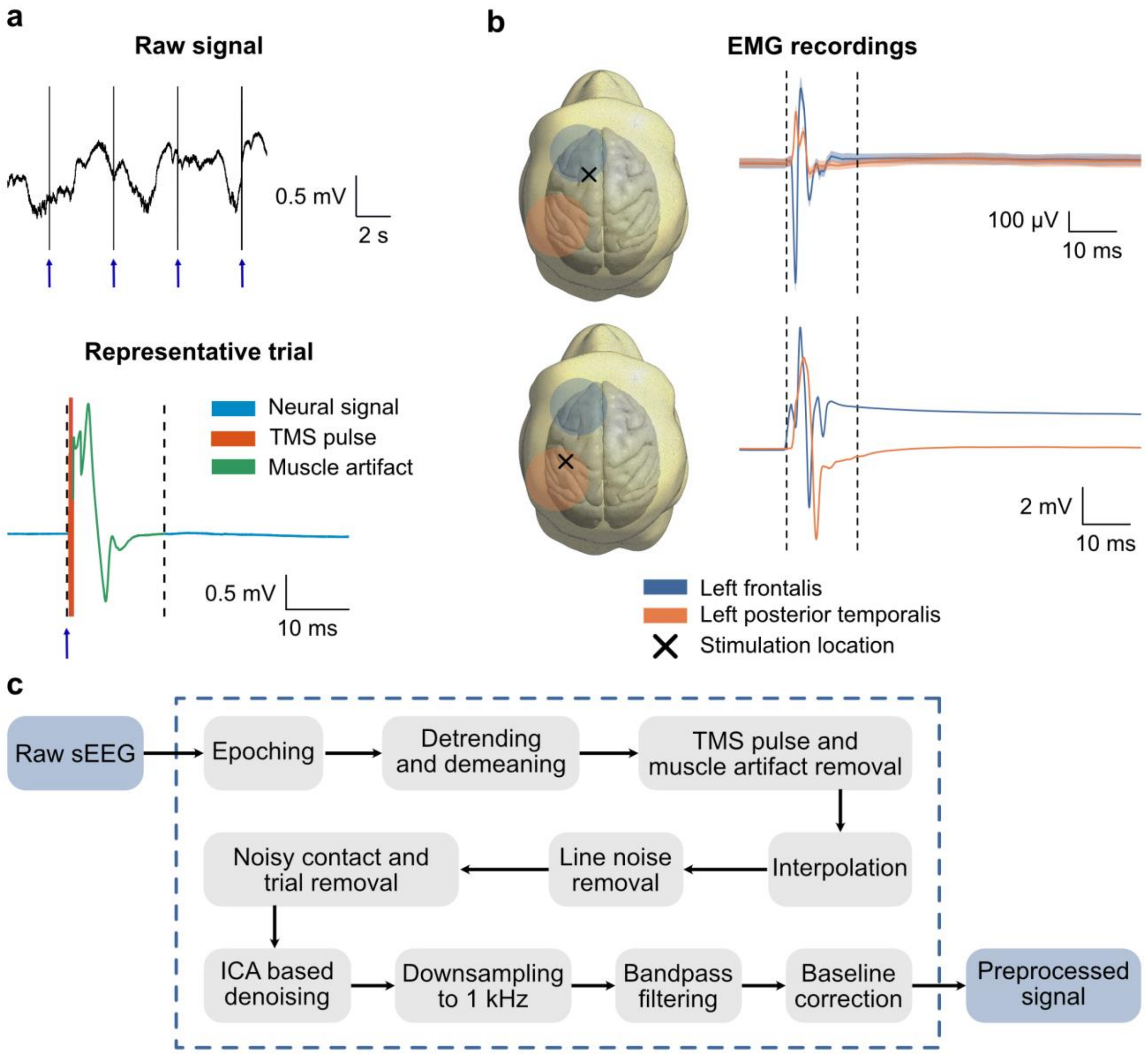


**Figure 1-1. Preprocessing of the LFP data.** a) Top. A fragment of the raw sEEG signal with TMS pulses marked by arrows. Bottom. A representative trial depicting the presence of TMS pulse, muscle artifact and the neural signal in the sEEG recording. The range of amplitude has been truncated to clearly show the magnitude of the muscle artifact. b) Electromyographic (EMG) recordings obtained by needle electrodes inserted into left frontalis and left posterior temporalis muscles in Monkey W. The general area of the muscle is shaded on the reconstructed head model (blue – left frontalis, orange – left posterior temporalis). Top. Time-locked EMG signals resulting from stimulating frontal brain region. Frontal muscle, that is closest to the stimulation site shows higher EMG activity compared to posterior muscle. Bottom. Time-locked EEG signals resulting from stimulating posterior brain region. In both cases, the muscle activity does not exceed the 15 ms time window that is shown in dashed vertical lines. Stimulation locations are shown by X. Time-locked EEG signals are calculated by interpolating the TMS pulse duration and calculating the average across trials. c) The detailed pipeline used for preprocessing sEEG signals.

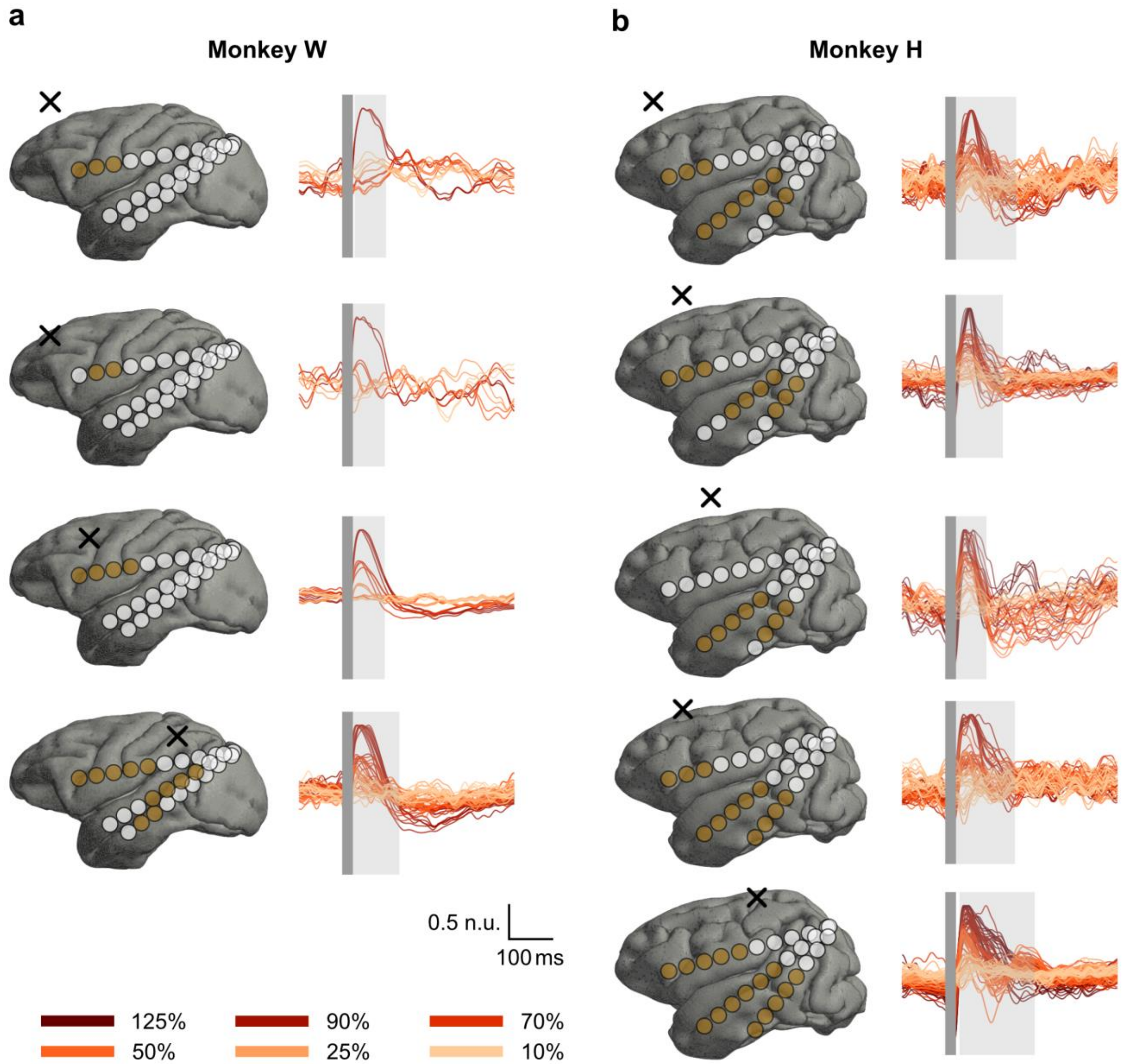


**Figure 2-1. Local field potentials from responsive contacts for each stimulation location.** a) Time-locked averaged signal from responsive contacts for each main stimulation location (1-4) in Monkey W. b) Time-locked averaged signals from responsive contacts for each main stimulation location (1-5) in monkey H. All the signals are normalized to the maximum amplitude of each contact. The responsive contacts are shaded in orange on the 3D rendering of the brain. The time window during which a significant effect of stimulation intensity on the LFPs is observed, is shaded in gray for each stimulation location.

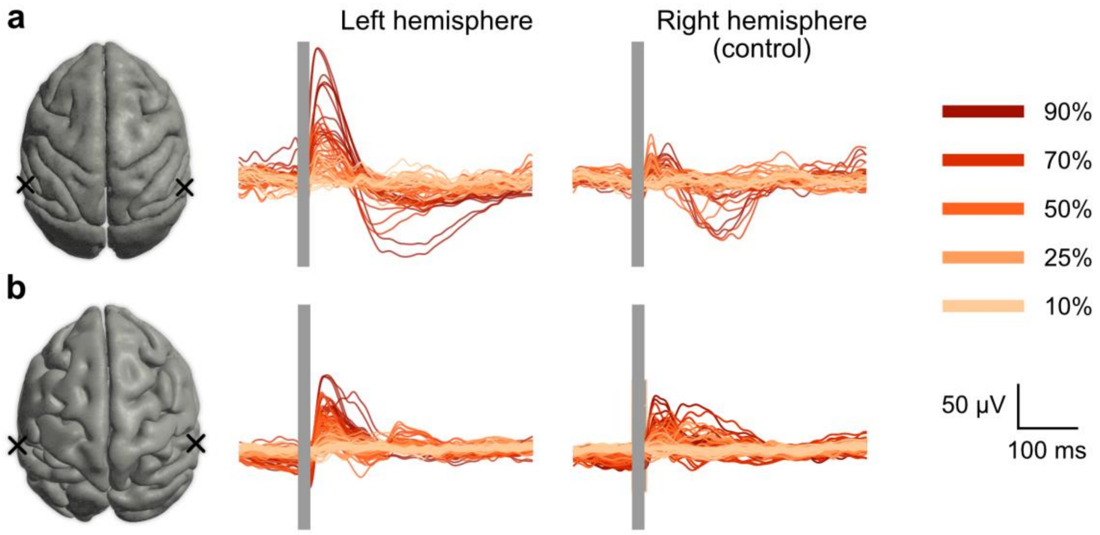


**Figure 2-2. Comparison of local field potentials between left and right hemispheres.** a) The butterfly plot of the local field potentials from all channels when stimulating location 4 (ipsilateral to recording hemisphere) and location 5 (contralateral control location) in Monkey W. In Monkey W, the overall activity is lower when stimulating the control location compared to stimulation location 4. b) Same plots for Monkey H (location 5 – ipsilateral to recording hemisphere and location 6 – contralateral control location). In Monkey H, the control location induces activity in the recording hemisphere albeit less than that of location 5.

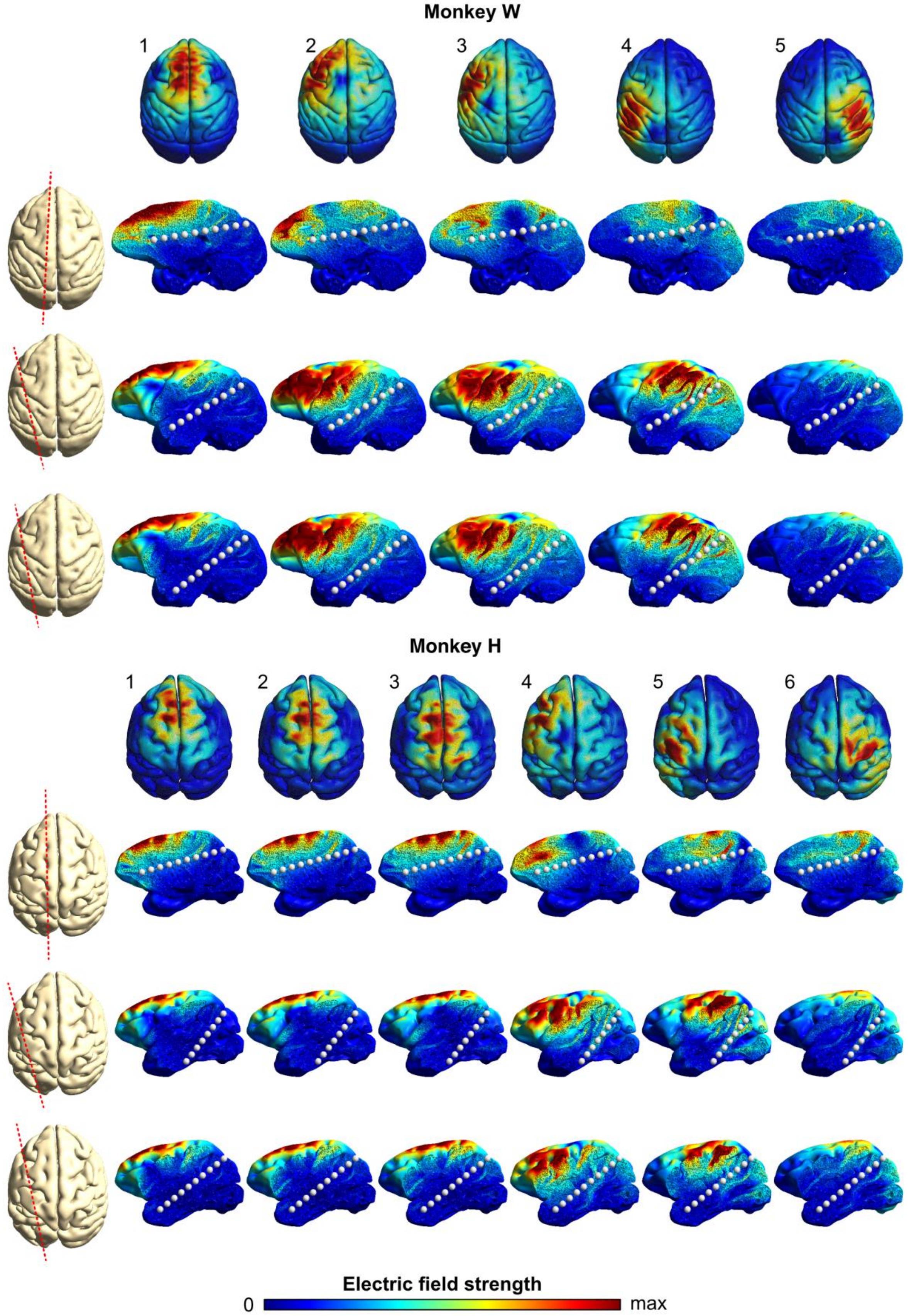

Monkey W
1
2
3
4
5
Monkey H
1
2
3
4
5
6
Electric field strength
0
max

**Figure 3-1. Profile of the penetration of electric fields at electrode locations.** The simulation results show the electric field magnitude captured by the electrode sites. The cross sections along the depth electrodes are shown in dotted red lines and the white dots show the contacts of the electrodes. The maximum is the 75th percentile of the electric field magnitude. We observe that significant electric fields reach the frontal regions in almost all stimulation locations which would potentially contribute to TEPs. However, electric fields rarely penetrate into the lateral regions, and the TEP responses from these regions could be a result of activation due to neural circuits.

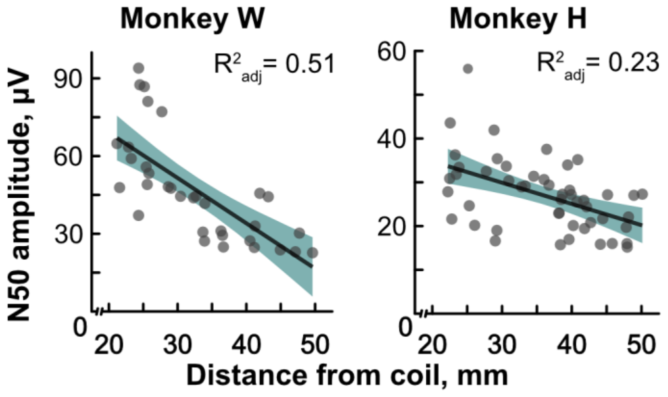


**Figure 4-1: Relationship between N50 response and the coil-to-contact distance.** Scatter plots show the amplitude of the N50 response against the coil-to-contact distance for each stimulation location. Stimulation locations 1-3 from monkey W and 1-4 from monkey H are included in this analysis. Linear regression analysis revealed a negative linear relationship between the N50 amplitude and the coil-to-contact distance in both monkeys W ($r = -0.81$, $p < 10^{-4}$) and H ($r = -0.52$, $p < 10^{-3}$).

**Table 1-1. Monkey W – The anatomical regions associated with each contact of the sEEG electrodes.** The anatomical regions were extracted using the post-implantation MR image, referenced to the common stereotaxic atlas. The endpoints of the depth electrodes were frontal eye field (FEF), auditory cortex (AUD) and temporal cortex (TEM). Abbreviations: (CN – caudate nucleus, CC – corpus callosum, AC – anterior cingulate, PCC – posterior cingulate cortex, MIP – medial intraparietal area, LIP – lateral intraparietal area, PO – preoptic area, S2 – secondary somatosensory cortex, STP – superior temporal plane, MT – middle temporal area, V4 – visual area 4, V3 – visual area 3, V2 – visual area 2, FST – fundus of superior temporal visual area, OFC – orbitofrontal cortex, MST – medial superior temporal area

| Contacts | FEF | AUD | TEM |
|---|---|---|---|
| 1 | CN (Head) | Auditory belt | TE |
| 2 | CN | Auditory belt/Insula | TE/STP |
| 3 | CN | Auditory belt/Insula | STP |
| 4 | CN | Auditory core/Insula | FST/STP |
| 5 | AC/CC | S2 | FST/STP |
| 6 | Dorsal bank of PCC (Area c)/CC | STP | MT |
| 7 | MIP | MT/STP | FST/TEO |
| 8 | MIP/LIP | V4 | V4 |
| 9 | PO/V3 | V2 | V2 |

**Table 1-2. Monkey H – The anatomical regions associated with each contact of the sEEG electrodes.** The same method used for Monkey W was to identify the anatomical regions. The endpoints of the depth electrodes were frontal eye field (FEF), auditory cortex (AUD) and inferior temporal region (IT). Abbreviations: (CN – caudate nucleus, OFC – orbitofrontal cortex, CC – corpus callosum, AC – anterior cingulate, MIP – medial intraparietal area, LIP – lateral intraparietal area, S2 – secondary somatosensory cortex, MT – middle temporal area, V3 – visual area 3, V2 – visual area 2, V1 – visual area 1, FST – fundus of superior temporal visual area, dPCG – dorsal posterior cingulate gyrus, MST – medial superior temporal area

| Contacts | FEF | AUD | IT |
|---|---|---|---|
| 1 | CN/OFC | Ventral insular | Parahippocampal |
| 2 | CN | Ventral insular | FST |
| 3 | CN/CC | Temporal/Insular cortex | FST |
| 4 | CN/CC/AC | Insular cortex/Auditory lateral belt | FST/MT |
| 5 | CN/CC/AC | Auditory core/Insular cortex | MT |
| 6 | CC | Auditory core/S2 | MT |
| 7 | CC | Auditory belt | WM |
| 8 | dPCG /MIP/LIP | Auditory belt/Brodmann's area 7A | WM |
| 9 | MIP | Brodmann's area 7A | MT |
| 10 | V1/V2 | V3/V2 | MT/MST |

**Table 2-1. Monkey W – Detailed statistics of cluster-based permutation tests to identify main effects of intensity.** The table shows the time window during which a significant main effect of stimulation intensity on LFP amplitude was observed, the maximum F-statistic in the given time window and the p-value. To identify the main effect of intensity, a permutation test was conducted. The F-statistic was calculated for each time point and was clustered based on temporal adjacency. Then the cluster statistic was calculated as the sum of the F-statistics. Finally, a permutation distribution of cluster statistics was generated with 1000 random permutations and the p-value was calculated by comparing the observed cluster statistic against the permutation distribution. For each stimulation location, the statistics of the responsive contacts are given here.

| | FEF | AUD | TEM |
|---|---|---|---|
| Location 1 | 29-101 ms, $F_{max}$ = 9.59, p = 0.003<br>29-102 ms, $F_{max}$ = 11.89, p = 0.003<br>30-96 ms, $F_{max}$ = 7.93, p = 0.037 | No significant contacts | No significant contacts |
| Location 2 | 25-99 ms, $F_{max}$ = 9.53, p = 0.005<br>25-82 ms, $F_{max}$ = 6.45, p = 0.046 | No significant contacts | No significant contacts |
| Location 3 | 25-83 ms, $F_{max}$ = 20.06, p = $9.9{\times}10^{-4}$<br>25-96 ms, $F_{max}$ = 24.16, p = $9.9{\times}10^{-4}$<br>25-99 ms, $F_{max}$ = 18.84, p = $9.9{\times}10^{-4}$<br>26-94 ms, $F_{max}$ = 12.36, p = 0.0017 | No significant contacts | No significant contacts |
| Location 4 | 25-133 ms, $F_{max}$ = 38.71, p = $9.9{\times}10^{-4}$<br>25-133 ms, $F_{max}$ = 49.52, p = $9.9{\times}10^{-4}$<br>25-117 ms, $F_{max}$ = 47.38, p = $9.9{\times}10^{-4}$<br>25-108 ms, $F_{max}$ = 33.17, p = $9.9{\times}10^{-4}$<br>25-96 ms, $F_{max}$ = 13.27, p = $9.9{\times}10^{-4}$ | 25-81 ms, $F_{max}$ = 19.93, p = 0.004<br>25-91 ms, $F_{max}$ = 23.92, p = $9.9{\times}10^{-4}$ | 25-110 ms, 1 $F_{max}$ = 21.65, p = $9.9{\times}10^{-4}$<br>25-99 ms, $F_{max}$ = 13.60, p = 0.003<br>25-96 ms, $F_{max}$ = 11.69, p = 0.01<br>25-97 ms, $F_{max}$ = 9.35, p = 0.02 |

**Table 2-2. Monkey W – Detailed statistics of cluster-based permutation tests to identify main effects of intensity.** The same procedure as for Monkey was carried out to identify the main effect of stimulation intensity on the LFP amplitude.

| | FEF | AUD | IT |
|---|---|---|---|
| Location 1 | 25-88 ms, $F_{max} = 10.52$, $p = 0.02$<br>35-80 ms, $F_{max} = 6.63$, $p = 0.05$<br>25-93 ms, $F_{max} = 12.95$, $p = 0.0015$ | 37-165 ms, $F_{max} = 4.40$, $p = 0.013$<br>31-161 ms, $F_{max} = 6.23$, $p = 0.006$<br>32-127 ms, $F_{max} = 7.78$, $p = 0.001$<br>29-128 ms, $F_{max} = 8.21$, $p = 9.9\times10^{-4}$<br>27-126 ms, $F_{max} = 14.20$, $p = 9.9\times10^{-4}$<br>28-125 ms, $F_{max} = 10.37$, $9.9\times10^{-4}$ | 25-86 ms, $F_{max} = 5.46$, $p = 0.041$<br>25-90 ms, $F_{max} = 6.17$, $p = 0.023$ |
| Location 2 | 37-93 ms, $F_{max} = 7.39$, $p = 0.03$<br>31-89 ms, $F_{max} = 11.47$, $p = 0.003$<br>27-90 ms, $F_{max} = 23.31$, $p = 9.9\times10^{-4}$ | 25-100 ms, $F_{max} = 8.08$, $p = 0.02$<br>25-95 ms, $F_{max} = 8.72$, $p = 0.019$<br>35-134 ms, $F_{max} = 9.64$, $p = 0.003$<br>34-129 ms, $F_{max} = 6.64$, $p = 0.01$ | 35-83 ms, $F_{max} = 8.41$, $p = 0.014$<br>32-92 ms, $F_{max} = 8.61$, $p = 0.09$<br>33-86 ms, $F_{max} = 9.02$, $p = 0.014$ |
| Location 3 | No significant contacts | 35-112 ms, $F_{max} = 5.08$, $p = 0.045$<br>32-107 ms, $F_{max} = 7.73$, $p = 0.015$<br>25-102 ms, $F_{max} = 7.51$, $p = 9.9\times10^{-4}$<br>25-78 ms, $F_{max} = 9.17$, $p = 0.045$<br>31-140 ms, $F_{max} = 12.42$, $p = 9.9\times10^{-4}$ | 25-93 ms, $F_{max} = 6.24$, $p = 0.022$<br>25-96 ms, $F_{max} = 9.19$, $p = 0.003$<br>25-86 ms, $F_{max} = 11.21$, $p = 0.003$ |
| Location 4 | 32-95 ms, $F_{max} = 8.08$, $p = 0.042$<br>30-119 ms, $F_{max} = 9.90$, $p = 0.001$<br>25-130 ms, $F_{max} = 18.25$, $p = 9.9\times10^{-4}$ | 31-143 ms, $F_{max} = 6.98$, $p = 0.07$<br>25-125 ms, $F_{max} = 7.90$, $p = 0.013$<br>31-90 ms, $F_{max} = 8.47$, $p = 0.036$<br>34-85 ms, $F_{max} = 18.17$, $p = 0.05$<br>32-92 ms, $F_{max} = 8.87$, $p = 0.025$ | 33-111 ms, $F_{max} = 5.31$, $p = 0.03$<br>33-117 ms, $F_{max} = 6.60$, $p = 0.015$<br>33-92 ms, $F_{max} = 11.79$, $p = 0.012$<br>35-113 ms, $F_{max} = 10.48$, $p = 0.007$ |
| Location 5 | 30-100 ms, $F = 25.22$, $p = 9.9\times10^{-4}$<br>29-133 ms, $F = 31.80$, $p = 9.9\times10^{-4}$<br>25-133 ms, $F = 44.03$, $p = 9.9\times10^{-4}$<br>25-132 ms, $F = 24.04$, $p = 9.9\times10^{-4}$<br>36-125 ms, $F = 12.17$, $p = 9.9\times10^{-4}$ | 34-141 ms, $F = 18.85$, $p = 9.9\times10^{-4}$<br>34-140 ms, $F = 15.53$, $p = 9.9\times10^{-4}$<br>34-145 ms, $F = 20.40$, $p = 9.9\times10^{-4}$<br>33-143 ms, $F = 22.61$, $p = 9.9\times10^{-4}$<br>33-138 ms, $F = 19.51$, $p = 9.9\times10^{-4}$<br>35-128 ms, $F = 10.99$, $p = 0.01$ | 33-174 ms, $F = 31.64$, $p = 9.9\times10^{-4}$<br>34-143 ms, $F = 17.78$, $p = 9.9\times10^{-4}$<br>35-157 ms, $F = 13.88$, $p = 9.9\times10^{-4}$<br>35-160 ms, $F = 13.36$, $p = 9.9\times10^{-4}$<br>37-162 ms, $F = 10.03$, $p = 9.9\times10^{-4}$ |
